# Nucleotide-pair encoding of 16S rRNA sequences for host phenotype and biomarker detection

**DOI:** 10.1101/334722

**Authors:** Ehsaneddin Asgari, Philipp C. Münch, Till R. Lesker, Alice C. McHardy, Mohammad R.K. Mofrad

**Affiliations:** Department of Bioengineering, University of California, Berkeley, CA 94720, USA; Computational Biology of Infection Research, Helmholtz Centre for Infection Research, Brunswick 38124, Germany; Max von Pettenkofer-Institute of Hygiene and Medical Microbiology, Faculty of Medicine, LMU Munich, 80336 Munich, Germany; Molecular Biophysics and Integrated Bioimaging, Lawrence Berkeley National Lab, Berkeley, CA 94720, USA

## Abstract

Identifying combinations of taxa distinctive for microbiome-associated diseases is considered key to the establishment of diagnosis and therapy options in precision medicine and imposes high demands on accuracy of microbiome analysis techniques. We propose subsequence based 16S rRNA data analysis, as a new paradigm for microbiome phenotype classification and biomarker detection. This method and software called DiTaxa substitutes standard OTU-clustering or sequence-level analysis by segmenting 16S rRNA reads into the most frequent variable-length subsequences. These subsequences are then used as data representation for downstream phenotype prediction, biomarker detection and taxonomic analysis. Our proposed sequence segmentation called nucleotide-pair encoding (NPE) is an unsupervised data-driven segmentation inspired by Byte-pair encoding, a data compression algorithm. The identified subsequences represent commonly occurring sequence portions, which we found to be distinctive for taxa at varying evolutionary distances and highly informative for predicting host phenotypes. We compared the performance of DiTaxa to the state-of-the-art methods in disease phenotype prediction and biomarker detection, using human-associated 16S rRNA samples for periodontal disease, rheumatoid arthritis and inflammatory bowel diseases, as well as a synthetic benchmark dataset. DiTaxa identified 17 out of 29 taxa with confirmed links to periodontitis (recall= 0.59), relative to 3 out of 29 taxa (recall= 0.10) by the state-of-the-art method. On synthetic benchmark data, DiTaxa obtained full precision and recall in biomarker detection, compared to 0.91 and 0.90, respectively. In addition, machine-learning classifiers trained to predict host disease phenotypes based on the NPE representation performed competitively to the state-of-the art using OTUs or k-mers. For the rheumatoid arthritis dataset, DiTaxa substantially outperformed OTU features with a macro-F1 score of 0.76 compared to 0.65. Due to the alignment- and reference free nature, DiTaxa can efficiently run on large datasets. The full analysis of a large 16S rRNA dataset of 1359 samples required ≈1.5 hours on 20 cores, while the standard pipeline needed ≈6.5 hours in the same setting.

**Availability:** An implementation of our method called DiTaxa is available under the Apache 2 licence at http://llp.berkeley.edu/ditaxa.

## 1 Introduction

Microbial communities vary widely in their taxonomic structures and compositions [1, 2, 3]. The human microbiota fulfills important functions in supporting, regulating, and causing adverse conditions in their environment, motivating methods for inferring relationships between microbial taxa or functions associated with certain host phenotypes. Due to its low cost, a popular data type generated in microbiome studies, is 16S rRNA amplicon data. The 16S rRNA gene includes both variable and conserved regions and is universally present in archaeal and bacterial microorganisms [4, 5, 6]. Particular regions of the 16S rRNA gene are amplified from degenerate primers and sequenced. After sequencing, reads are typically clustered based on their sequence similarity to each other and the resulting clusters are referred to as operational taxonomic units (OTUs). Three main strategies for creating OTUs have been developed: in the *de novo* OTU clustering scheme, input sequences are aligned against one another and OTU clusters created based on a user-specified percent identity cutoff (in practice mostly 97%) without comparisons to reference databases. The implementation of the *de novo* strategy is difficult to parallelize and therefore limited to small-scale datasets. Variations of this method, such as sub-sample open-reference OTU picking [7] or centroid-based greedy clustering approaches [8] accelerate this process and enable their application to larger datasets. Alternatively, in closed-reference OTU clustering, input reads are aligned to a set of cluster centroids defined in a reference database (containing clusters of previously identified OTUs) and will be reported as an OTU, if they align at a given threshold. This strategy will not report OTUs for novel taxa that are not part of the reference database, though. An advantage is the usual high quality of taxonomic assignments of the reference database, which can be used for taxonomic assignment of the OTUs from the community of interest. Finally, the open-reference OTU clustering scheme combines *de novo* and closed-reference picking, where input sequences are aligned against a reference database (such as Greengenes [9]) and sequences that fail to match the reference are subsequently clustered *de novo* in a serial process [7]). Individual algorithms for OTU clustering, post- and pre-processing have been combined to pipelines such as mothur [10], QIIME [11, 12], USEARCH [13] and LotuS [14].

Although OTU clustering has simplified 16S rRNA processing by substituting the analysis of millions of reads by analysis of only thousands of OTUs, it still has several disadvantages: OTUs do not necessarily represent meaningful taxonomic units, such as e.g. species, and sequencing errors may inflate diversity estimates by orders of magnitude [15]. To prevent diversity overestimates, OTU based approaches require a highly stringent quality control and relaxed clustering at < 97% similarity. While this approach limits the inflation of OTUs by potential sequencing errors, it comes at the expense of taxonomic resolution and may combine organisms with distinct biological properties and capabilities into a single OTU. A further disadvantage is that OTU calling requires extensive sequence alignment efforts. All of the above mentioned OTU-picking strategies involve sequence alignments either to the reference genomes or to the sample sequences, which is computationally expensive and cannot be easily extended to further samples. It was shown that OTUs were generally ecologically consistent across habitats, but observed OTU content can differ substantially between clustering methods [16]. Since the number of obtained OTUs and their content is dependent on the pipeline and the parameter settings, reproducing the same analysis is difficult [17]. An alternative solution is the analysis of individual 16S rRNA gene sequence [18, 19, 20], which is computationally challenging, as each 16S rRNA sample may contain 10,000s of sequences.

Popular machine learning tasks over 16S rRNA gene sequencing data are taxonomic classification, host phenotype prediction, as well as biomarker detection. Although k-mer features and some other non-OTU features have been also used [18, 21], the most common representation of 16S rRNA gene sequences is based on OTUs. Random Forest was reported as the most effective classification approach for several diseases [2, 3, 21, 22, 23]. Recently, we have shown that using k-mer representations of shallow-subsamples is computationally inexpensive (being reference- and alignment-free) and marginally outperforms OTUs in host phenotype and environment classification tasks [21]. However, a disadvantage of k-mer features is that short k-mers cannot easily be mapped to a taxonomy to obtain taxonomic biomarkers. Microbiome studies often aim to identify OTUs, taxa, or clades that differ in their abundance across two or more subsets of the input samples (e.g. between diseased and healthy states), here referred as biomarker discovery [24, 25]. Identifying these biological informative taxa that are enriched in only a subset of phenotypes (e.g. diseased subjects or patients that better respond to a certain treatment) is a challenging task, in particular for metagenomic samples, because of their high-dimensionality, sequencing errors, as well as other systematic biases, such as the presence of chimeric sequences [15]. One prominent biomarker example is the over-representation of the Firmicutes phylum in obese individuals compared to lean controls [26, 27, 28]. In case the over-representations are causal for the aetiology of the diseases, detection of such biomarkers might have potential therapeutic implications if disease progression can be reversed by targeting over-expressed causative species using emerging technologies such as CRISPR/Cas9 or phage-based targeting [29, 30, 31]. This is also true for biomarkers that are inversely related to disease progression, such as the over-representation of *Akkermansia muciniphila* in individuals with a healthier metabolic status and better clinical outcome after caloric restriction [32]. For many other diseases where a microbiological component is expected, such (combinations) of biomarkers yet have to be found. Even when the biomarkers fail to be causal, they may enable prediction of the disease state or disease sub-types, or suggest suitable therapies in personalized medical interventions.

Different methods have been developed to identify OTU-based biomarkers [33]. The most widely used method is linear discriminant analysis effect size (LEfSe), which has a particular focus on high-dimensional class comparison for metagenomic analysis and determines features (such as taxa, OTUs, genes or clades) most likely to explain differences between two or more classes from relative OTU abundances [34]. This method uses the non-parametric factorial Kruskal-Wallis (KW) sum rank test [35]. Several other with similar functionality exist that use different statistical tests over sample profiles based on OTU features, such as STAMP [36], MetaStats [34] and MetagenomeSeq [33]).

In this paper, we propose DiTaxa, an alignment- and reference-free, subsequence based paradigm for processing of 16S rRNA microbiome data for phenotype and biomarker detection. DiTaxa substitutes standard OTU-clustering by segmenting 16S rRNA sequences into variable length subsequences. The obtained subsequences are then used as data representation for downstream phenotype and biomarker detection. We show that DiTaxa outperforms the state-of-the-art approach in biomarker detection for synthetic and a number of disease-related datasets. In addition, DiTaxa performs competitively with the k-mer based state-of-the-art approach, outperforming OTU-features, in phenotype prediction.

## 2 Material and Methods

### 2.1 Datasets

#### Inflammatory Bowel Diseases

We use the largest pediatric Crohn’s disease dataset available to date, described in [37]^1^, which covers different types of Inflammatory Bowel Diseases (IBD). This is a dataset of 1359 labeled 16S rRNA samples from 731 pediatric (≤ 17 years old) patients diagnosed with Crohn’s disease (CD), 219 with ulcerative colitis (UC), 73 with indeterminate colitis (IC), and 336 samples verified as healthy. Sequencing was targeted towards the V4 hypervariable region of the 16S rRNA gene. We downloaded OTU representations of the samples from Qiita repository^2^ obtained using QIIME pipeline [7].

#### Rheumatoid arthritis

We downloaded read data (454 platform) of the 16S rRNA gene sequences of V1 and V2 rRNA for 114 fecal DNA samples of a rheumatoid arthritis (RA) study [38] from SRA (ID: SRP023463). OTU clustering was performed based on filtered reads (365.7*k*, 23.0%) of which 140,382 were unique and 119,217 singletons and resulted in 949 OTUs based on 97% identity. The OTU clustering pipeline is detailed in §2.3.

#### Periodontal disease

We use the data provided by Jorth, *et al*. [39] to differentiate between healthy and diseased periodontal microbiota^3^. This dataset consists of microbial samples collected from subgingival plaques from 10 healthy and 10 patients diagnosed with periodontitis. Sequencing was targeted towards the (*V*4 − *V*5) hypervariable region of the 16S rRNA gene. Similar to the RA dataset, we obtain the OTU features using the clustering pipeline detailed in §2.3.

#### Synthetic dataset

To evaluate DiTaxa in a known setting, we generated a dataset with synthetic samples using Grinder v. 0. 5.3 [40] based on 1000 V4 regions of different genera of Green-genes (GG) sequences [9]. V4 regions were extracted from the Green-genes 13.8 databases using the forward and reverse primer sequences GTGCCAGC[AC]GCCGCGGTAA and ATTAGA[AT]ACCC[CGT][AGT]GTAGTCC. To generate 16S rRNA datasets, the *length_b_ias* parameter was set to zero and the *unidirectional* parameter was set to one. To cover the full V4 region, the amplicon read length distribution was set to 300 and the fold coverage of the input reference sequence was set to 30. We specified the percent of reads in the amplicon libraries that should be chimeric sequences to 10%. We used default parameters for the specification of the chimera distribution resulting in 89% bimeras, 11% trimeras and 0.3% quadmeras. Sequencing errors were introduced in the reads, at positions that follow a uniform model using the default ratio of substitutions to the number of indels (4 substitutions for each indel). Two sets of samples were created, denoted as *case* and *control* samples with an average number of sequences in both groups of 29,204 reads. While in the control set all 1000 genera were set to the same abundance (mean abundance set to 0.1%, 500 randomly selected GG V4 sequences (corresponding to unique genera) were enriched at equal levels in the case dataset (μ of non-selected genera set to 0.05%; *μ* of selected genera set to 0.15;). For both, the control and the case settings, 100 samples were generated, each with variations under the normal distribution (*σ* = 0.02). We processed the synthetic dataset using a standard pipeline consisting of USEARCH and UPARSE and generated 1,041 OTUs at 97% identity similarity, detailed in §2.3.

### 2.2 Nucleotide-pair Encoding

The idea of Nucleotide-pair Encoding (NPE) is inspired by the Byte Pair Encoding (BPE) algorithm, a simple universal text compression scheme [41, 42], which has been also used for compressed pattern matching in genomics [43]. Although BPE had lost its popularity for a long time in compression, only recently it again became popular, but for a different reason, i.e. word segmentation in machine translation in natural language processing (NLP). BPE became a common approach for a data-driven unsupervised segmentation of words into their frequent subwords, which facilitate open vocabulary neural network machine translation and improve the quality of translation by reducing the vocabulary size [44, 45]. In this work, we adapt the BPE algorithm for splitting biological sequences into frequent variable length subsequences called Nucleotide-pair Encoding (NPE). We propose NPE as general purpose segmentation for the biological sequence (DNA, RNA, and proteins). In contrast to the use of BPE in NLP for vocabulary size reduction, we use this method to increase the size of symbols from 4 nucleotides to a large set of variable length biomarkers.

The input to NPE is a set of sequences. We treat each sequence as a list of characters (nucleotides in the case of 16S rRNA gene sequences). The algorithm finds the most frequently occurring pair of adjacent symbols in the sequences. On the next, we replace all instances of the selected pair with a new subsequence (merged pair as a new symbol). The algorithm repeats this process until reaching a certain vocabulary size or when no more frequently occurring pairs of symbols available. The obtained merging operations can be inferred once from a large set of sequences in an offline manner and then applied to an unseen set of sequences. A simple pseudo-code of NPE is provided in Algorithm 1.

#### Algorithm 1 Adapted Byte-pair algorithm (BPE) for segmentation of biological sequences (NPE)

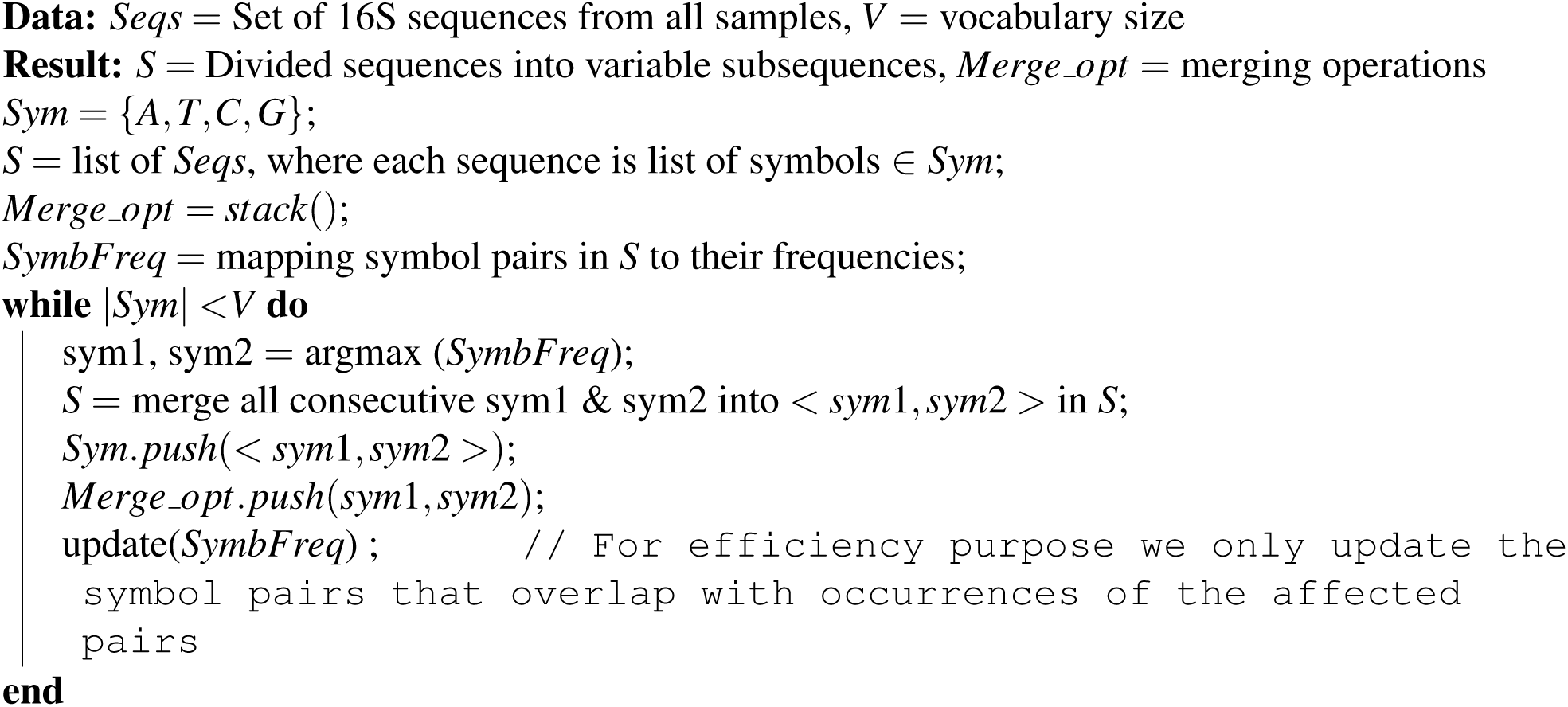

### 2.3 Standard 16S rRNA gene processing workflow

To evaluate the performance of DiTaxa against the state-of-the-art, we used a standard 16S rRNA gene processing workflow employed in previous studies on 16S rRNA data [46, 47, 48]. Note that throughout this paper “the standard pipeline (STDP)” refers to this workflow:

1. Obtained 16S rRNA gene sequencing reads are quality controlled and clustered using the Usearch 8.1 software package^4^, where quality filtering is done with *fastq_filter*(*–fastq_maxee*1).
2. The OTU clusters and representative sequences are determined using the UPARSE algorithm (*derep_fulllength* : *minuniquesize*2;*cluster_otus* : *otu_radius_pct*3) [49].
3. The next step is taxonomy assignment using the EZtaxon database [50] as the reference database, and the decision is made by RDP Classifier [51].
4. The OTU absolute abundance table and mapping file are used for statistical analyses in LDA Effect Size (LEfSe) [34].

### 2.4 DiTaxa computational workflow

The DiTaxa computational workflow has three main components; (i) NPE representation creation, (ii) phenotype prediction, and (iii) biomarker detection and taxonomic analysis (shown in Figure 1). In this section, these components are described in details.

**Figure 1.**
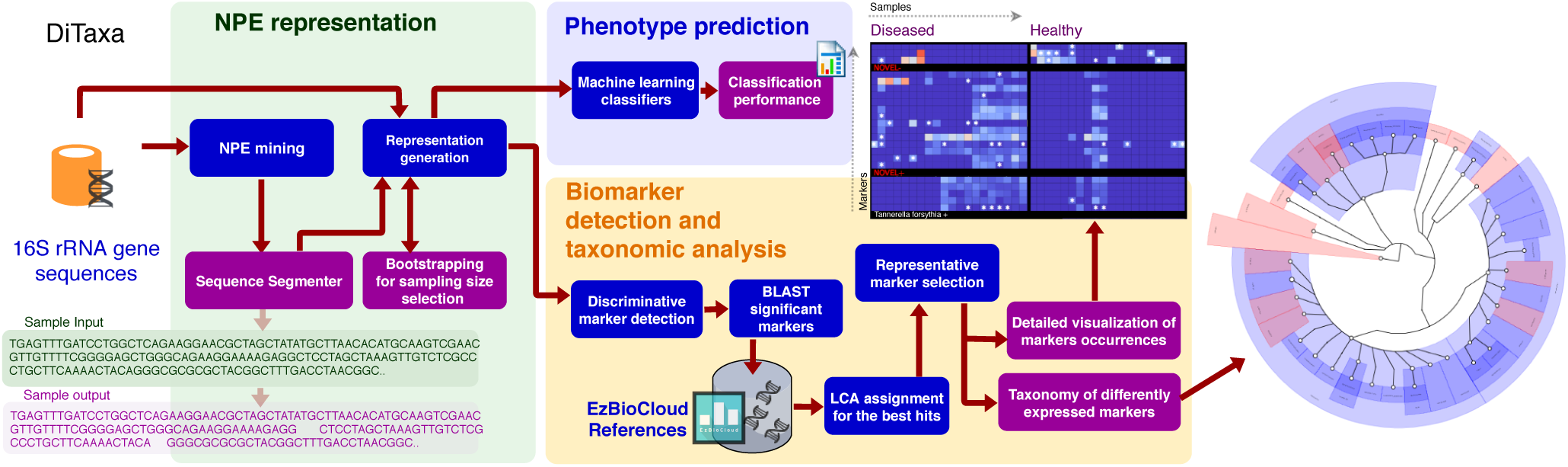
Computational workflow of DiTaxa, DiTaxa has three main components: (i) NPE representation, (ii) Phenotype prediction, (ii) Biomarker detection and taxonomic analysis. The purple boxes denote the outputs of the approach.

#### NPE representation

The first component of DiTaxa is the NPE representation creation. The 16S rRNA gene sequences aggregated from all samples from all phenotypes go through the NPE algorithm 1 for training segmentation operations. Then the segmentation will be applied on sequences to segment sequences into variable length subsequences. We pick the vocabulary size large enough to obtain discriminative 16S rRNA subsequences considered as biomarkers. Each sample will be presented as a count distribution of its subsequences. We propose a bootstrapping scheme to investigate the sufficiency of shallow sub-samples to produce proper representation.

In a previous study, using a bootstrapping framework we showed that shallow sub-samples of 16S rRNA gene sequences are sufficient to produce a proper k-mer presentation of data for phenotype prediction [21]. Similarly, here we use bootstrapping to investigate sufficiency and consistency of NPE representation, when only a small portion of the sequences are used. This has two important implications, first, sub-sampling reduces the preprocessing run-time, second, it shows that even a shallow 16S rRNA sequencing is enough for the phenotype prediction. We use a resampling framework to find a proper sampling size. Let θ*_#npe_*(*X_i_*) be the normalized NPE (with vocabulary size of #*npe*) distribution of *X_i_*, a set of sequences in the *i^th^* 16S rRNA sample. We investigate whether only a portion of *X_i_*, which we represent as 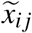, i.e. *j^th^* resample of *X_i_* with sample size N, would be sufficient for producing a proper representation of *X_i_*. To find a sufficient sample size for *X_i_* quantitatively, we propose the following formulation in a resampling scheme. **(i) Self-consistency**: resamples for a given size *N* from *X_i_* produce consistent 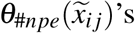, i.e. resamples should have similar representations.(ii) Representativeness: resamples for a given size *N* from *X_i_* produce 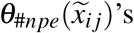 similar to *θ_#npe_*(*X_i_*), i.e. similar to the case where all sequences are used. As presented in [21], we measure the **self-inconsistency** 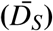 of the resamples’ representations by calculating the average Kullback Leibler divergence among normalized NPE distributions for *N_R_* resamples (here *N_R_*=10) with sequences of size *N* from the *i^th^* 16S rRNA sample:

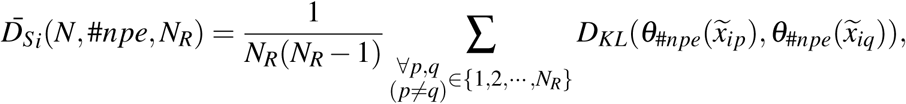

where 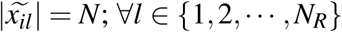. We calculate the average of the values of 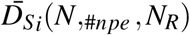 over the *M* different 16S rRNA samples:

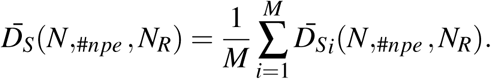

We measure the **unrepresentativeness** 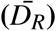 of the resamples by calculating the average Kullback Leibler divergence between normalized NPE distributions for *N_R_* resamples (*N_R_*=10) with size *N* and using all the sequences in *X_i_* for the *i^th^* 16S rRNA sample:

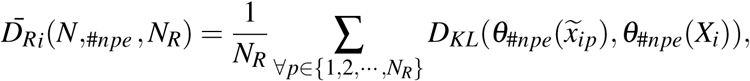

where 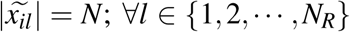. We calculate the average over 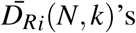 for the *M* 16S rRNA samples:

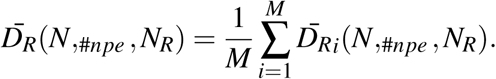

For the experiments on the datasets presented in §2.1, we measure self-inconsistency 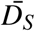 and unrepresen-tativeness 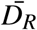 for *N_R_* = 10 and *M* = 10 for #*npe* ∈ {10000,20000,50000} with sampling sizes ranging from 20 to 20000.

As shown in Figure 1, the obtained NPE representation in the first component will be then used for two main use cases, i.e. phenotype prediction and biomarker detection.

#### Phenotype prediction

We used Random Forest (RF) classifiers [52], which have shown a superior performance over deep neural network (deep multi-layer perceptron) and support vector machine (SVM) classifiers in phenotype classification for the size of datasets we use here [21, 22]. However, the provided implementation provides deep learning and SVM classifiers as well. For the disease phenotype prediction, Random Forest classifiers were tuned for (i) the number of decision trees in the ensemble, (ii) the number of features for computing the best node split, and (iii) the function to measure the quality of a split. We evaluate and tune the model parameter using stratified 10 fold cross-validation and optimize the classifiers for the harmonic mean of precision and recall, i.e. the F1-score, as a trade-off between precision and recall. We provide both micro- and macro-F1 metrics, which are averaged over instances and over categories, respectively.

We performed phenotype classification for a synthetic dataset (binary classification of 100 case samples and 100 control samples), a Crohn’s disease dataset (binary classification of 731 Crohn’s disease samples from 628 control or other diseases), and a Rheumatoid Arthritis (RA) dataset (44 RA disease subjects versus 70 control/treated/Psoriatic arthritis subjects). In order to evaluate the performance of NPE representation, we compare the classification performance of RFs over NPE features versus using OTUs, as well as k-mer features, which are considered as state-of-the-art approaches for disease phenotype prediction [21, 22].

#### Biomarker detection and taxonomic analysis

The designed steps in DiTaxa for detection of differently expressed markers in the phenotype of interest are shown in the light purple background in Figure 1:

1. The first step is finding discriminative markers between two phenotype states using false discovery rate corrected two-sided *χ*^2^ test over the median-adjusted presence of markers in the samples. Thus if a marker is presented within a sample at least as frequent as the median frequency across samples, we consider it as present, otherwise as absent. We discard insignificant markers using a threshold for the p-value of < 0.05. For the multi-phenotype case, a one-versus-all policy is used. In addition, markers shorter than a certain threshold (< 50*bps*) will be discarded to ensure the markers are specific enough for a downstream taxonomic assignment.
2. The filtered markers go through a local BLAST [53] with EzBioCloud database as a local reference dataset [50], covering 62,362 quality controlled reference sequences. We assign the taxon corresponding to the Lowest Common Ancestor (LCA) of the taxa annotated for the best hits of a marker in a reference taxonomy. The markers that cannot be aligned to the references will be marked as ‘Novel’ markers.
3. In the third step, we remove redundant markers based on their co-occurrence information using symmetric Kullback-Leibler divergence [54]:

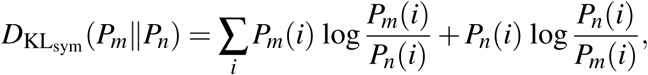

where *P_m_* and *P_n_* are respectively normalized frequency distributions of *m^th^* and *n^th^* markers across all samples. Using 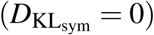 to find identical markers, split the set of markers into equivalence classes. Subsequently, from each class we pick only one representative marker with the most specific taxonomy level, which its taxonomy information is confirmed by the majority of markers within the class. The selected markers at this step are our final set of biomarkers.
4. Our approach has three main outputs, first, a taxonomic tree for significant discriminative biomarkers, where identified taxa to the positive and negative class are colored according to their phenotype (red for positive class and blue for negative class). The DiTaxa implementation for taxonomic tree generation uses a Phylophlan-based backend [55]. Second, a heatmap of top biomarkers occurrences in samples, where the rows denote markers and the columns are samples is generated. Such a heatmap allows biologists to obtain a detailed overview of markers’ occurrences across samples (e.g., Figure 10 and Figure 11). The heatmap shows number of distinctive sequences hit by each biomarker in different samples and stars in the heatmap denote hitting unique sequences, which cannot be analyzed by OTU clustering approaches. The third output is a list of novel markers for further analysis of the potential novel organisms.

To compare the performance of our approach with a standard workflow (defined in §2.3) for real datasets, we used the scientific literature as the ground-truth. We extracted a list of organisms which are experimentally identified by previous studies to be associated with or cause periodontal disease. Then we evaluate the recall for DiTaxa and the standard workflow in the detection of the confirmed organisms.

To quantify the performance of DiTaxa in a known synthetic setting versus the standard pipeline, we generated two high-dimensional synthetic datasets denoted as ‘case’ and ‘control’, as described in §2.1. The description of the standard pipeline is provided in §2.3 generating a list of significant differently expressed OTUs for both phenotypes. We then compare the significantly enriched OTU sequences (FDR corrected P values < 0.05, LEfSe) and significant (FDR < 0.05) subsequences determined by DiTaxa as biomarkers with the ground-truth 16S V4 region using global nucleotide alignment with blastn v. 2.7.1+ with parameters “–*perc_identity* 100 – *ungapped*”. We quantified the number of false positives (FP), true positives (TP), false negatives (FN) and true negatives (TN) based on the presence of significant alignments of potential biomarkers sequences (OTU or DiTaxa markers) to each of the 500 differentially expressed GG 16S regions. TPs were calculated as the number of GG sequences (*n* = 500) with at least one marker hit from the positive marker list. FNs were calculated using case GG sequences (*n* = 500) without at least one marker hit from the marker collection found to be significantly enriched in the case set. TNs are the number of control GG sequences (*n* = 500) with at least one marker hit from the marker list that are significant in the control set. FPs were quantified as the number of low abundant GG sequences (*n* = 500) without at least one marker hit from the set of markers found in the control sample. Recall was calculated as *TP*/(*TP* + *FN*) while precision was calculated as *TP*/(*TP* + *FP*).

## 3 Results

### 3.1 Phenotype prediction

#### Bootstrapping for sample size selection

We picked a stable sample size for each NPE vocabulary size based on the output of bootstrapping in phenotype prediction. Each point in Figure 2 represents the average of 100 (*M* × *N_R_*) resamples belonging to *M* randomly selected 16S rRNA samples, each of which is resampled *N_R_* = 10 times. As shown in Figure 2, a larger vocabulary size require higher sampling rates to produce self-consistent and representative representations. As the structure of the curve does not vary a lot from dataset to dataset, to avoid redundancy, we only show the bootstrapping results for the rheumatoid arthritis dataset.

**Figure 2.**
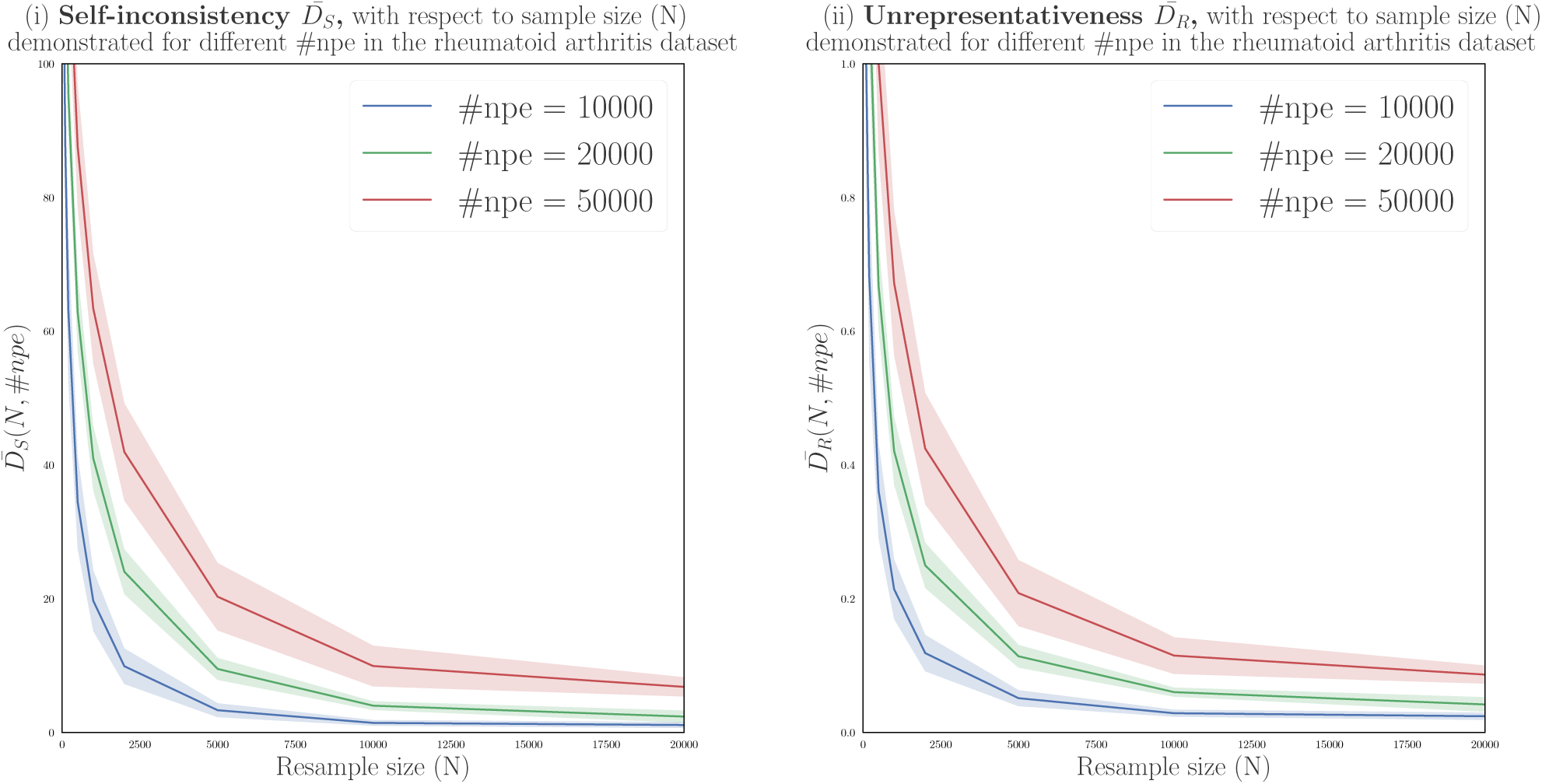
Measuring (i) self-inconsistency 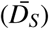, and unrepresentativeness 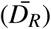 for the arthritis disease dataset. Each point represents an average of 100 resamples belonging to 10 randomly selected 16S rRNA samples. Higher vocabulary size require higher sampling rates to produce self-consistent and representative samples.

The classification results for different NPE vocabulary sizes on the synthetic, Crohn’s disease, and rheumatoid arthritis datasets using RF classifiers are presented in Table 1. All methods could reliably predict the affected cases in the synthetic dataset without any error. For the Crohn’s disease dataset k-mers with the MicroPheno approach [21] achieved a slightly better prediction performance while using NPE and OTU features achieved the same macro-F1 of 0.74. In rheumatoid arthritis prediction, NPE and k-mers achieved a macro-F1 of 0.76, outperforming the use of OTU features by 11 percent. Changes in sample size did not substantially affect the prediction performance, suggesting sufficiency of shallow sub-samples in phenotype prediction using the NPE representation (Table 1).

**Table 1.**
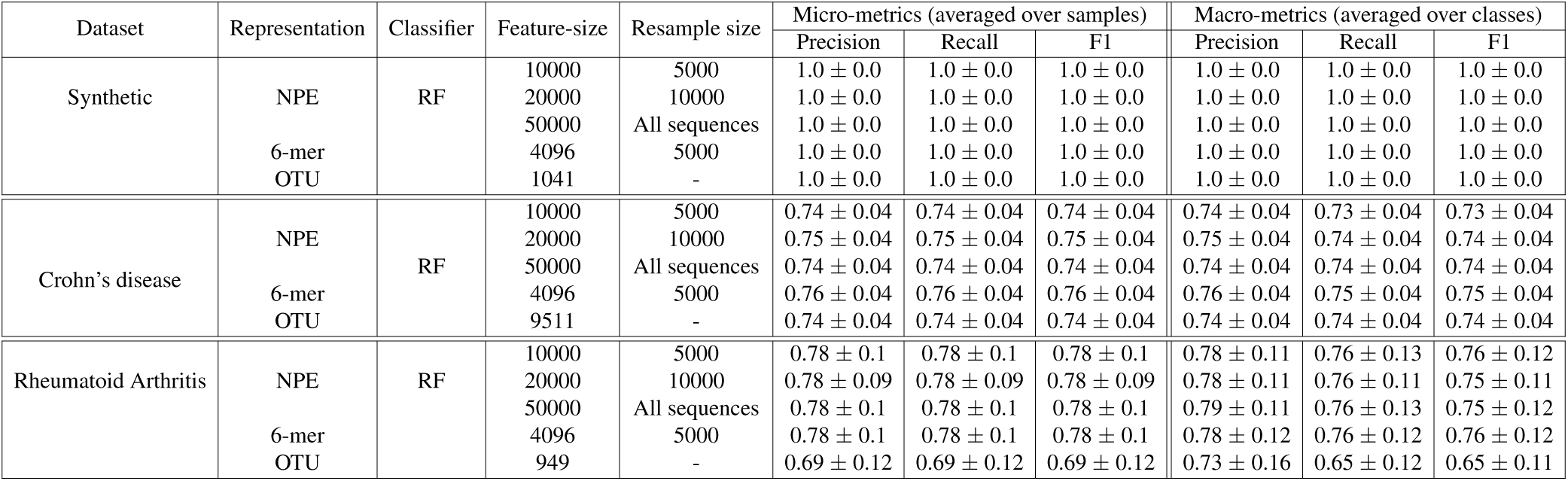
Results for NPEs, OTUs, and k-mer features in performing phenotype classification over the synthetic, rheumatoid arthritis, and Crohn’s disease datasets in a 10-fold cross-validation framework using random forest classifiers.

### 3.2 Biomarker detection and taxonomic analysis

#### Marker detection results for synthetic data

In the biomarker detection for the synthetic dataset, DiTaxa did not report erroneous (neither FN nor FP) biomarkers, which resulted in a recall and precision value of 1. In comparison, the biomarkers detected with a standard pipeline, using OTU clustering and LEfSe, included 51 false negative and 47 false positive instances, resulting in a recall and precision value of 0.898 and 0.905, respectively (Table 2). This evaluation demonstrated the superiority of DiTaxa for biomarker discovery compared to OTU-based approaches in both recall and precision.

**Table 2.** The results of DiTaxa and the standard pipeline (STDP) in marker detection for the synthetic dataset.

| Method | Precision | Recall | F1 |
| --- | --- | --- | --- |
| DiTaxa | 1 | 1 | 1 |
| STDP | 0.905 | 0.898 | 0.901 |

#### Biomarker detection results for periodontal disease

We next assessed the performance of DiTaxa and a standard pipeline (STDP; section 2.3) in detecting otherwise confirmed taxa for periodontal disease (Table 4). DiTaxa performed better in the detection of relevant taxa (Table 3). Of 29 taxa identified as relevant in other studies, 17 were detected by DiTaxa, while the standard approach detected only 3 from the same dataset. Notably, experimentally verified taxa shown to alter the disease phenotype in mouse models, *Fusobacterium nucleatum* [56] and *Porphyromonas gingivalis* [56, 57, 58], were only detected by Ditaxa. Since periodontitis is a polymicrobial disease and the oral biofilms are extremely diverse [59], detecting all relevant taxa confirmed by the literature from a single dataset is not feasible. For instance, A. actinomycetemcomitans is specifically associated with juvenile aggressive periodontists in Moroccan population, which can be hardly found in the population from Turkey [39]. However, relative comparison of recall for different methods on the same dataset is still meaningful. A higher recall of DiTaxa in confirming the literature links, in comparison with a standard pipeline shows that DiTqxa can be more accurate in detecting disease-specific biomarkers. A detailed comparison of the predicted taxa with different methods and taxa with confirmed links by the literature is also shown in Figure 3. The red color shows the disease associated taxa found by DiTaxa and the blue color indicates the up-regulated taxonomy in the healthy samples. The up-regulated organisms found by standard pipeline are colored in yellow and down-regulated organisms are colored to green. The intersection of DiTaxa and the standard approach is colored in orange for up-regulation and cyan denotes for the consensus of methods in down-regulation.

**Table 3.** The results of DiTaxa and the standard pipeline (STDP) in marker detection in comparison with literature of periodontal disease.

| Method | True Positive Count | Recall |
| --- | --- | --- |
| DiTaxa | 13 out of 29 | 0.59 |
| STDP | 3 out of 29 | 0.10 |

**Table 4.** Comparison of a standard pipeline (STDP) and DiTaxa performance in detecting 29 taxa with confirmed links to periodontitis. The upwards arrow denotes upregulation in the diesease group compared to samples from healthy subjects. A checkmark denotes presence of marker by DiTaxa method.

| Genus | Species | Direction | Literature information<br>Description and reference | Detection results |  |
| --- | --- | --- | --- | --- | --- |
|  |  |  |  | DiTaxa | STDP |
| Actinomyces | n.a. | ↓ | 16S rRNA sequencing of supragingival plaque samples, t-test [60], Periodontal disease vs. healthy control gingiva, t-tests [61] | not found | not found |
| Aggregatibacter | A. actinomycetemcomitans | ↑ | Used for mixed infections for murine back abscess model [62] | not found | not found |
| Anaeroglobus | n.a. | ↑ | Periodontal disease vs. healthy control gingiva, t-tests [61] | not found | not found |
| Corynebacterium | n.a. | ↓ | Periodontal disease vs. healthy control gingiva, t-tests [61] | not found | not found |
| Campylobacter | C. gracilis | ↑ | 16S rRNA sequencing [63] | ✓ | not found |
| Campylobacter | C. rectus | ↑ | 16S rRNA sequencing [63] | not found | not found |
| Campylobacter | C. showae | ↑ | 16S rRNA sequencing [63] | not found | not found |
| Dialister | n.a. | ↑ | Periodontal disease vs. healthy control gingiva, t-tests [61] | not found | not found |
| Eubacterium | E. nodatum | ↑ | 16S rRNA sequencing [63] | ✓ | not found |
| Fusobacterium | F. nucleatum | ↑ | Mouse model [56], Whole genomic DNA probes of subgingival plaque samples [64], and dot blot hybridization [65] | ✓ | not found |
| Fusobacterium | F. periodonticum | ↑ | microbial co-localization in images of subgingival biofilm species [66] | not found | not found |
| Gamella | n.a. | ↓ | 16S study of plaque samples from healthy individuals [67] | not found | not found |
| Granulicatella | n.a. | ↓ | 16S study of plaque samples from healthy individuals [67], Periodontal disease vs. healthy control gingiva, t-tests [61] | not found | not found |
| Leptotrichia | n.a. | ↑ | 16S rRNA sequencing of supragingival plaque samples, t-test [60] | ✓ | not found |
| Neisseria | n.a. | ↓, ↑ | Periodontal disease vs. healthy control gingiva, t-tests [61] (↓), Whole genomic DNA probes of subgingival plaque samples [68] (↑) | ✓ (↑) | not found |
| Parvimonas | P. micra | ↑ | 16S rRNA sequencing [59] | ✓ | not found |
| Porphyromonas | n.a. | ↑ | Mouse model [58] and Periodontal disease vs. healthy control gingiva, t-tests [61] | ✓ | not found |
| Porphyromonas | P. gingivalis | ↑ | Mouse model [56, 57], Whole genomic DNA probes of subgingival plaque samples [64], Used for monomicrobial infections for murine back abscess model [62] | ✓ | not found |
| Prevotella | n.a. | ↑ | 16S rRNA sequencing of supragingival plaque samples, t-test [60], Periodontal disease vs. healthy control gingiva, t-tests [61] | ✓ | not found |
| Prevotella | P. intermedia | ↑ | Whole genomic DNA probes of subgingival plaque samples [64] and dot blot hybridization [65] | ✓ | not found |
| Prevotella | P. nigrescens | ↑ | 16S rRNA sequencing [63] | not found | not found |
| Schwartzia | n.a. | ↑ | Periodontal disease vs. healthy control gingiva, t-tests [61] | ✓ | ✓ |
| Selenomonas | n.a. | ↑ | Periodontal disease vs. healthy control gingiva, t-tests [61] | ✓ | ✓ |
| Streptococcus | n.a. | ↓ | 16S rRNA sequencing of supragingival plaque samples, t-test [60], 16S study of plaque samples from healthy individuals [67] | ✓ | not found |
| Streptococcus | S. constellatus | ↑ | 16S rRNA sequencing of supragingival plaque samples, t-test [69] | ✓ | not found |
| Tannerella | T. forsythia | ↑ | Whole genomic DNA probes of subgingival plaque samples [64], 16S rRNA sequencing [70] | ✓ | not found |
| Treponema | n.a. | ↑ | Periodontal disease vs. healthy control gingiva, t-tests [61] | ✓ | ✓ |
| Treponema | T. denticola | ↑ | Whole genomic DNA probes of subgingival plaque samples [64], nested PCR [59] | ✓ | not found |
| Veillonella | n.a. | ↓ | 16S study of plaque samples from healthy individuals [67] | not found | not found |

**Figure 3.**
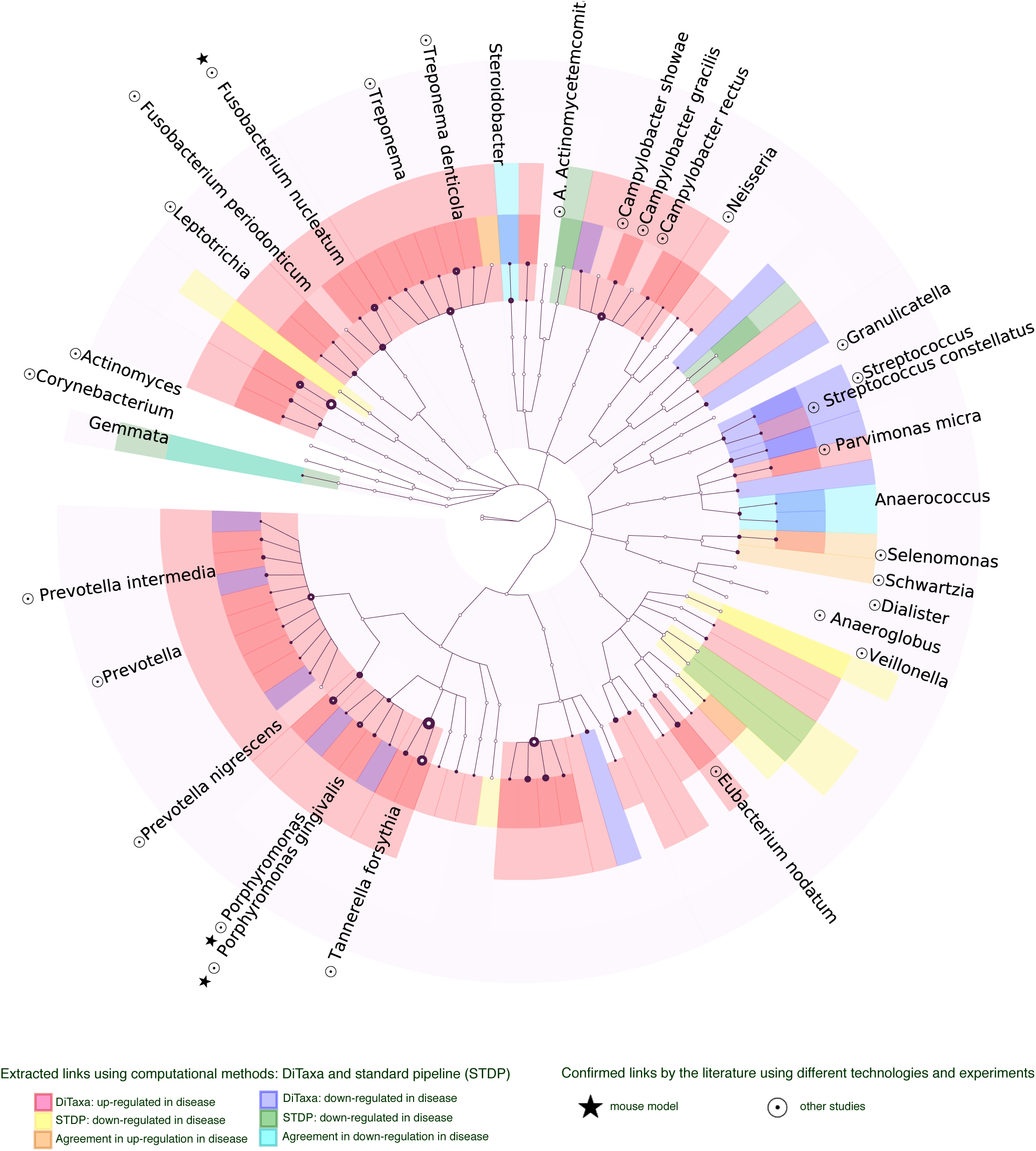
Taxonomy of differently expressed markers for samples from patients with periodontal disease versus healthy patients. The red color shows the disease associated taxa found by DiTaxa and the blue color indicates the up-regulated taxonomy in the healthy samples. The up-regulated organisms found by standard pipeline (STDP) are colored in yellow and down-regulated organisms are colored to green. The intersection of DiTaxa and the standard approach is colored in orange for up-regulation and cyan denotes for the consensus of methods in down-regulation. When two methods disagree the organism is colored in gray. The node size is proportional to the number of markers confirming the taxa and the border boldness is proportional to the negative log of the markers’ p-value. The taxa names are only provided for the nodes that are either confirmed by the literature or the nodes denoting agreements of DiTaxa and STDP.

#### Taxonomy of discriminative biomarkers for rheumatoid arthritis

Comparative taxonomic visualization of detected differentially expressed markers for DiTaxa and a common workflow are shown in Figure 4 for samples from patients with untreated rheumatoid arthritis (new onset RA) versus healthy individuals. Taxa predicted by DiTaxa for samples from patients with untreated rheumatoid arthritis (new onset RA) versus healthy individuals had *Prevotella copri* as the most significantly ranked, which was confirmed based on shotgun metagenome analysis and in mouse experiments [71], while the standard workflow only predicted the genus of this taxon as relevant [71]. DiTaxa also predicted *Prevotella stercorea* as implicated in new onset RA.

**Figure 4.**
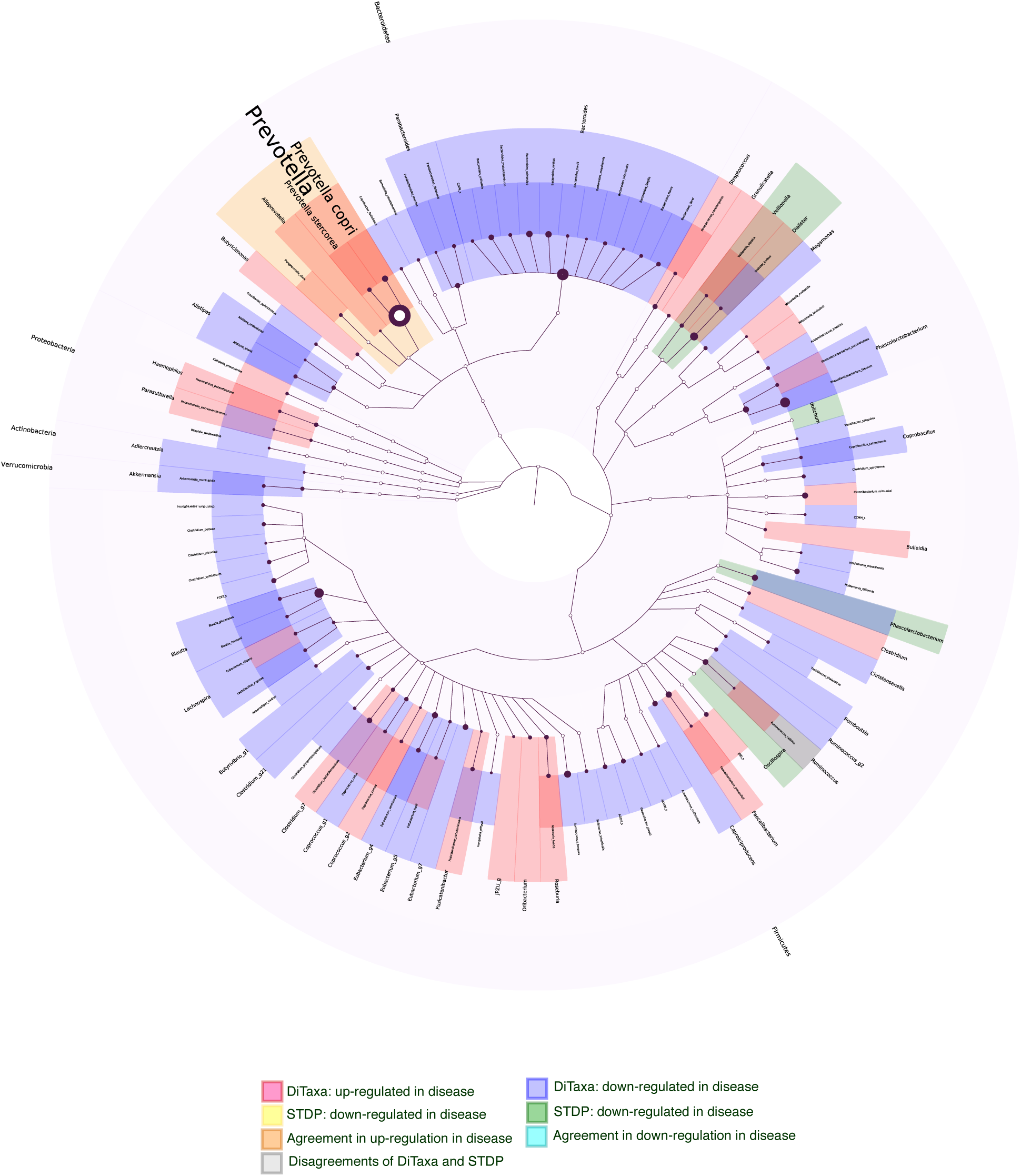
Taxonomy of differently expressed markers for new onset rheumatoid arthritis versus healthy samples. The red color shows the disease-associated taxa found by DiTaxa and the blue color denotes the up-regulated taxa in the samples from healthy individuals. The up-regulated organisms found by the standard pipeline are colored in yellow and down-regulated organisms are colored green. The agreements of DiTaxa and the standard approach are colored in orange for up-regulation and cyan denotes the consensus of both methods in down-regulated taxa. When the two methods disagree, the taxon is colored in gray. The node size is proportional to the number of markers confirming the taxa and the border boldness is proportional to the negative log of the markers’ p-value.

The DiTaxa results for patient samples from several other diseases versus healthy individuals are provided in Figure 5 (for CD versus healthy), Figure 7 (for indeterminate colitis versus healthy), Figure 6 (for ulcerative colitis versus healthy), Figure 9 (for treated rheumatoid arthritis versus healthy), Figure 8 (for psoriatic versus healthy).

**Figure 5.**
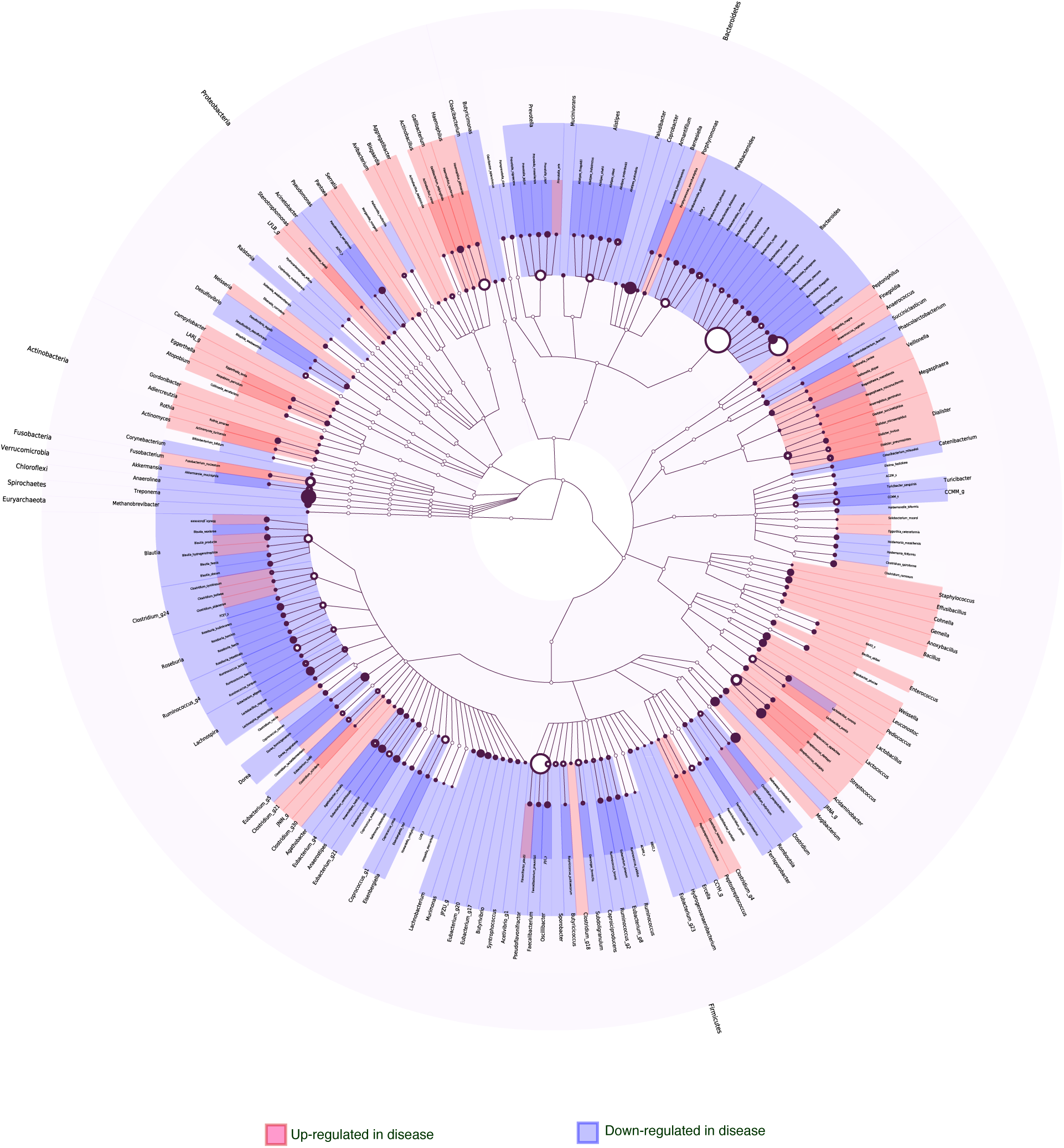
Taxonomy of differently expressed markers for Crohn’s disease versus healthy using DiTaxa. The red color shows the disease associated taxa found by DiTaxa and the blue color shows the up-regulated taxonomy in the healthy samples.

**Figure 6.**
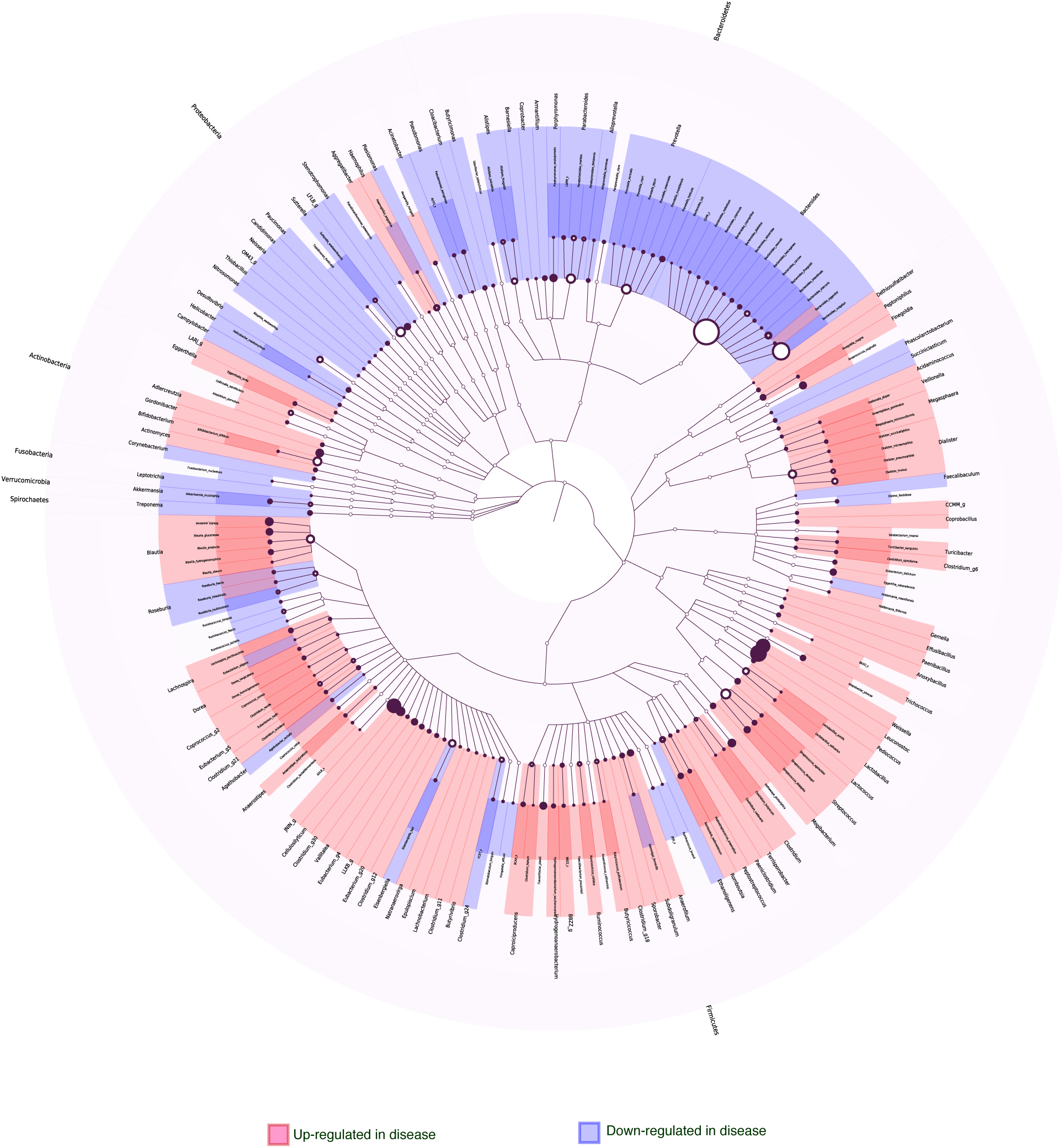
Taxonomy of differently expressed markers for ulcerative colitis disease versus healthy using DiTaxa. The red color shows the disease associated taxa found by DiTaxa and the blue color shows the up-regulated taxonomy in the healthy samples.

**Figure 7.**
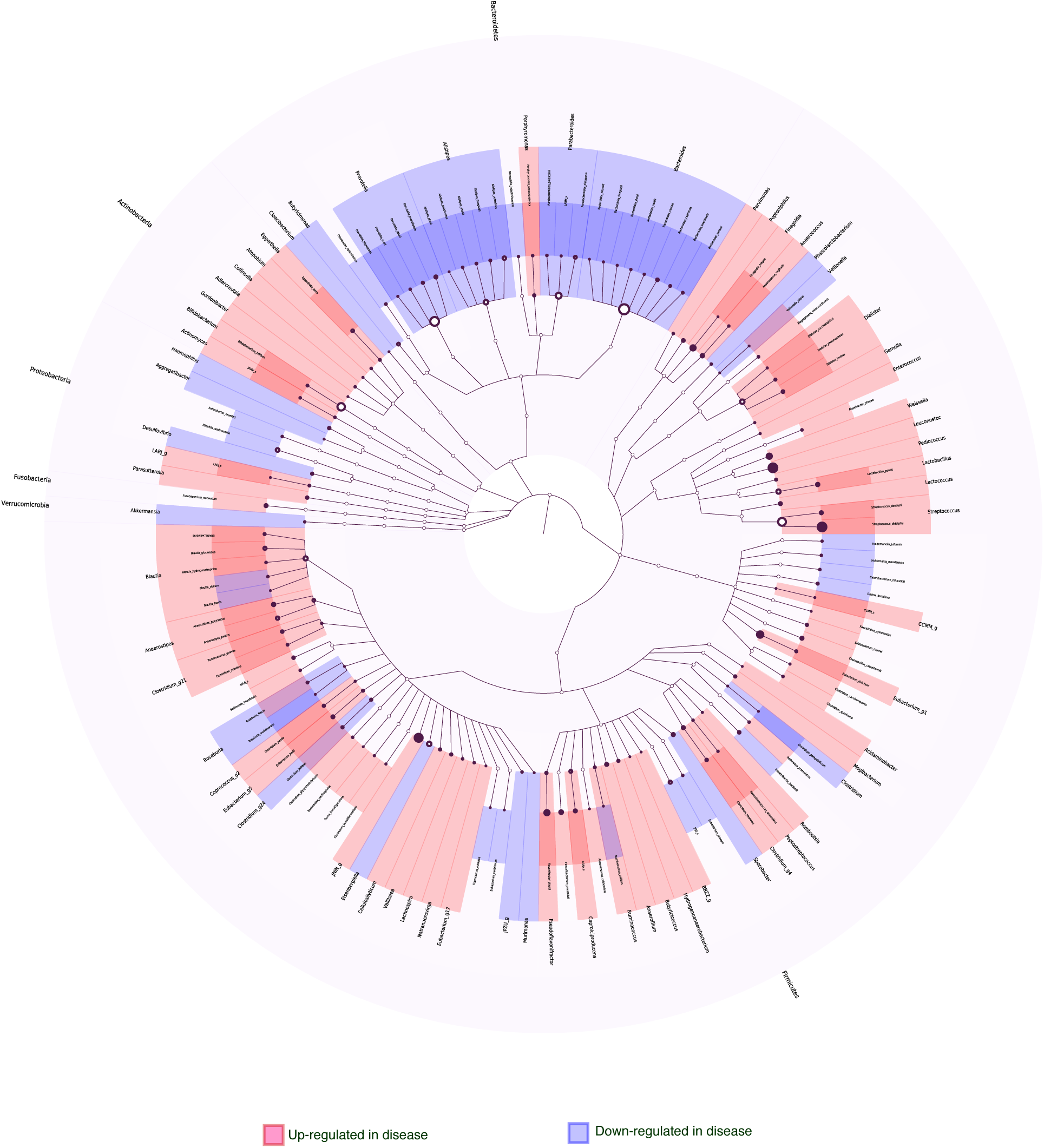
Taxonomy of differently expressed markers for indeterminate colitis disease versus healthy using DiTaxa. The red color shows the disease associated taxa found by DiTaxa and the blue color shows the up-regulated taxonomy in the healthy samples.

**Figure 8.**
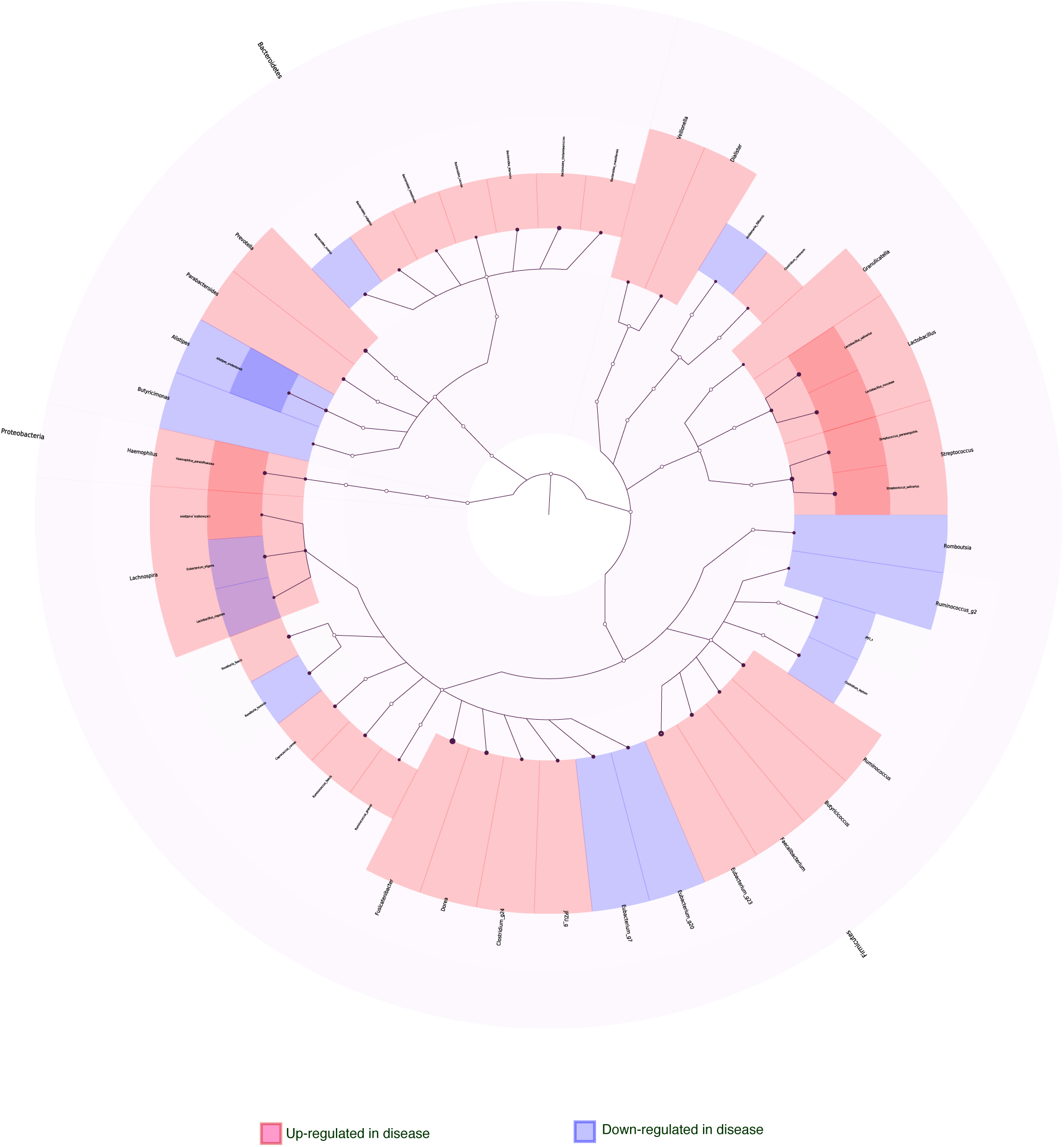
Taxonomy of differently expressed markers for psoriatric disease versus healthy using DiTaxa. The red color shows the disease associated taxa found by DiTaxa and the blue color shows the up-regulated taxonomy in the healthy samples.

**Figure 9.**
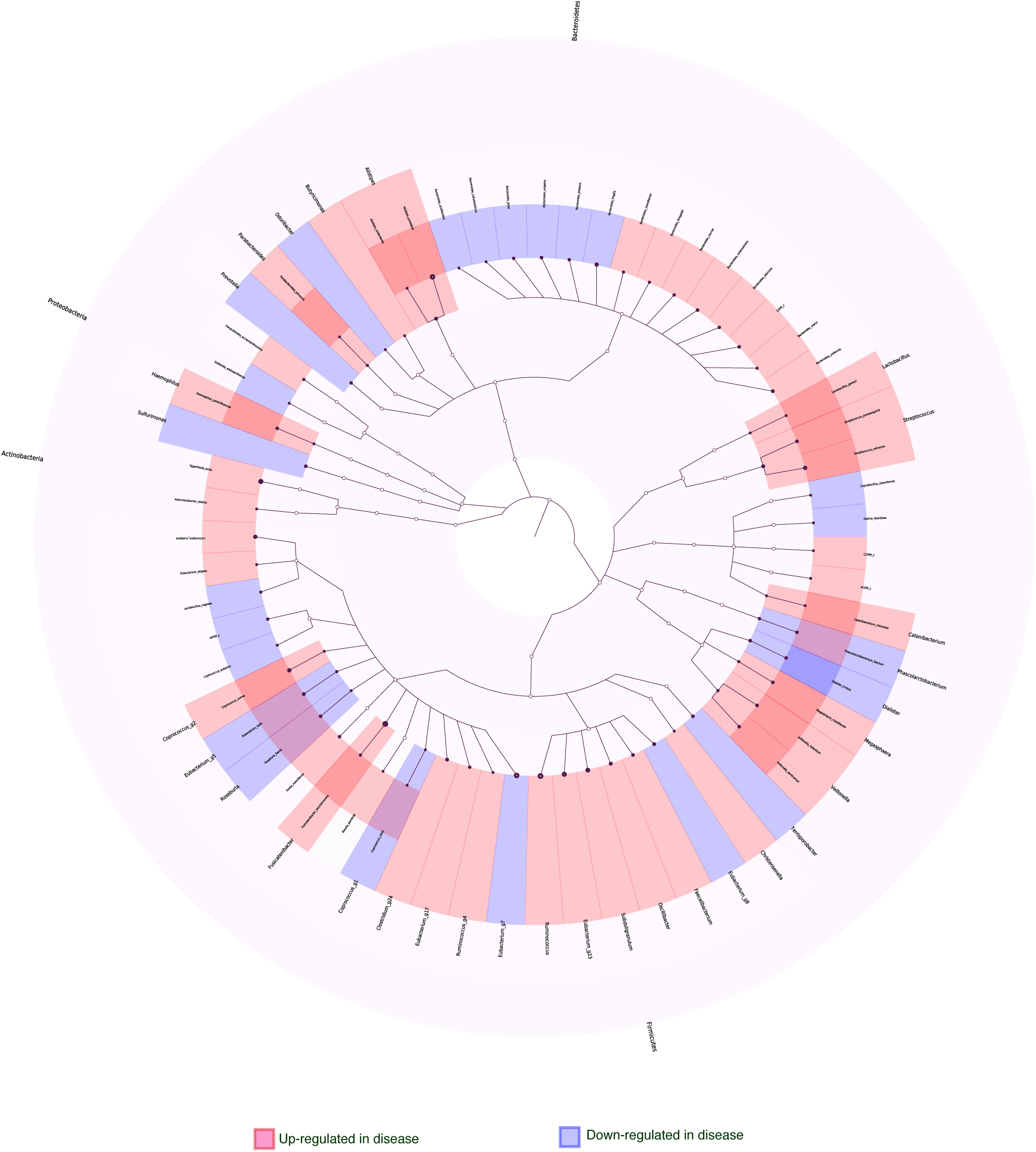
Taxonomy of differently expressed markers for treated rheumatoid Arthritis versus healthy samples using DiTaxa. The red color shows the treated associated taxa found by DiTaxa and the blue shows the up-regulated taxonomy in the healthy samples.

#### Biomarker heatmaps

Visualization of the occurrence pattern of the identified biomarkers is another output of DiTaxa. Examples of such a visualization for rheumatoid arthritis (Figure 10) and periodontal disease (Figure 11) are provided. The rows represent inferred biomarker sequences and are sorted based on the taxonomic marker assignments. The columns represent patient samples and are sorted firstly based on their phenotype and secondly based on their pattern similarity. ‘Novel’ organisms are shown in the top rows, denoting the markers that could not be aligned to any reference sequence and are therefore potentially novel taxa. The cell colors on the heatmap show the percentage of distinct marker sequences matching a biomarker per sample on a log scale. Markers targeting a single 16S sequence only are marked by a star.

**Figure 10.**
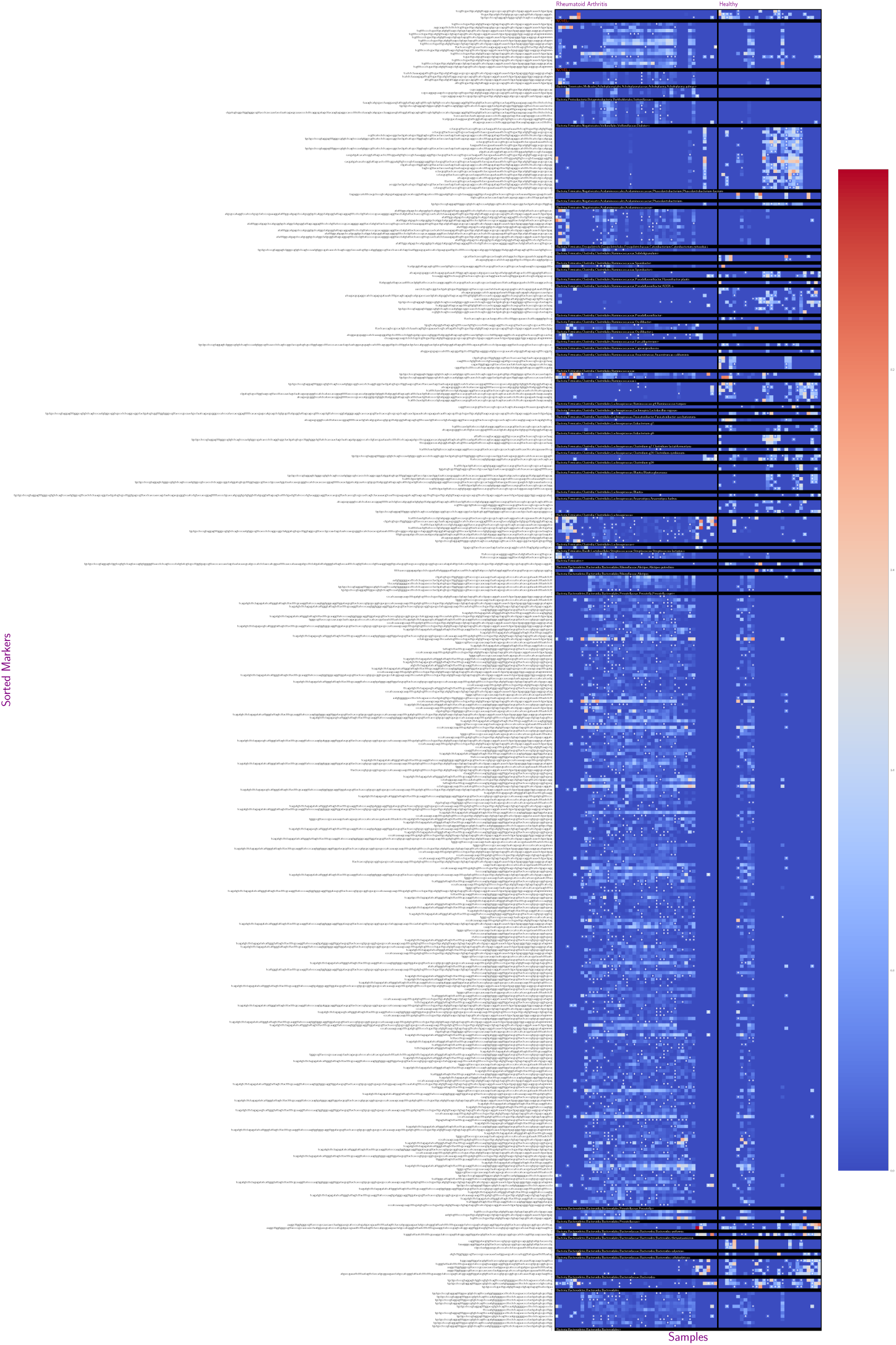
Heatmap of markers occurrences across samples for rheumatoid arthritis.

**Figure 11.**
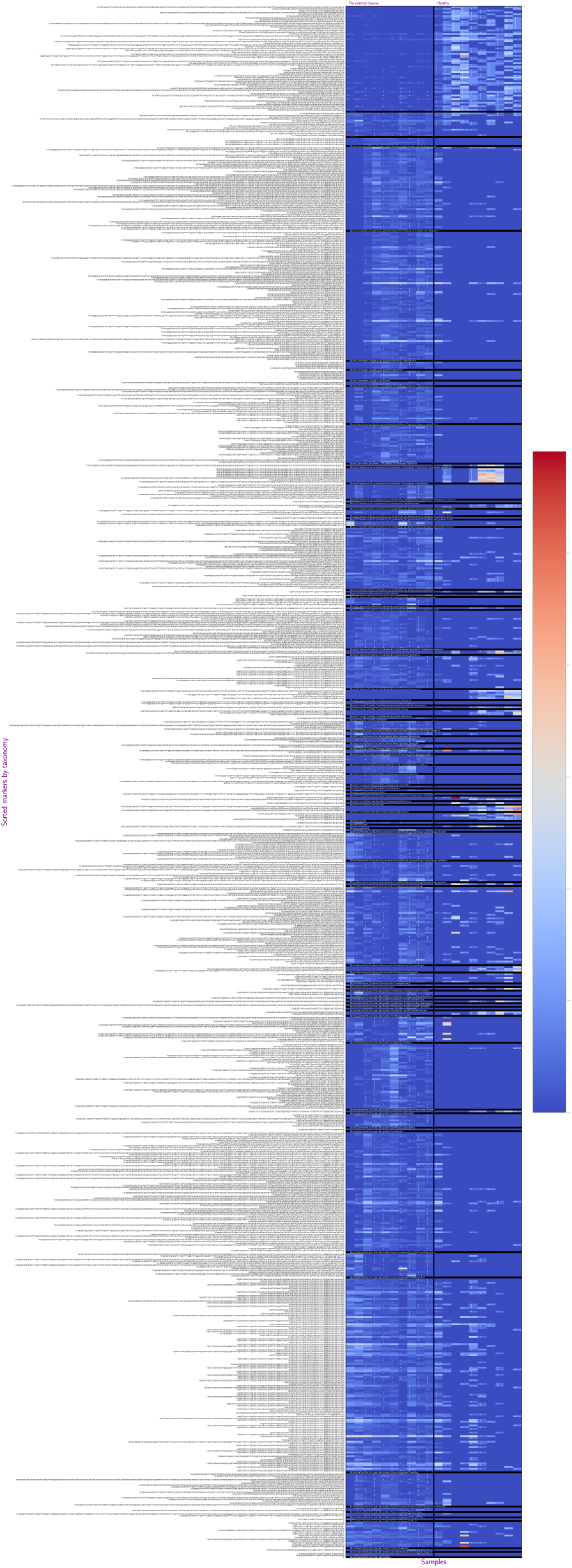
Heatmap of markers occurrences across samples for periodontal disease.

The plots clearly show the varying “generality” of the inferred marker sequences, with some matching only to unique 1S sequences, and others found across larger numbers, indicating representation of different levels of evolutionary relatedness of the underlying targeted organisms. For instance, of the inferred markers assigned *Prevotella copri*, while some markers match multiple distinct 16S genes across patient samples, indicating the presence of strain-level diversity evident from 16S within this species, while other markers targeting predominantly single 16S copies across patients samples within this species, indicating the existence of disease-associated subspecies diversity, that can be discovered with this technique.

### 3.3 Runtime analysis

To assess computational efficiency, we compared the runtimes of DiTaxa versus the standard workflow (Table 5). For both DiTaxa and the standard workflow 20 cores were used in computations. Workflow parts that could be parallelized for both pipelines are denoted with “||” in Table 5. The bottleneck for DiTaxa computation is the segmentation training, which cannot be parallelized. However, the segmentation needs to be trained only once for a dataset and then any combinations of phenotype analysis can use the trained segmentation and the subsequent representation. Although the standard pipeline for datasets of less than 200 samples has been few minutes faster than DiTaxa, DiTaxa can run faster for the dataset of 1359 samples (total of 93,93 min), while the standard pipeline tool 385,66 min using the same computational setting.

**Table 5.** Runtimes of DiTaxa and a common 16S processing pipeline for the periodontal, synthetic, rheumatoid arthritis and IBD datasets. The (||) sign denotes steps that are parallelized and run on multiple cores (here 20 cores). STDP stands for standard pipeline, using OTU clustering and LEfSe for marker detection.

| Dataset | Number of samples | Segmentation or clustering |  | Representation Creation |  | Biomarker detection |  | Total |  |
| --- | --- | --- | --- | --- | --- | --- | --- | --- | --- |
|  |  | DiTaxa | STDP (\|\|) | DiTaxa (\|\|) | STDP | DiTaxa(\|\|) | STDP (\|\|) | DiTaxa | STDP |
| Periodontal dataset | 20 | 3,37 min | 0,34 min | 1,84 min | 0,43 min | 1,58 min | 0,10 min | **6,79 min** | **0,87 min** |
| Rheumatoid Arthritis dataset | 114 | 23,86 min | 1,33 min | 0,61 min | 5,10 min | 1,73 min | 0,11 min | **26,19 min** | **6,54 min** |
| Synthetic dataset | 200 | 19 min | 0,81 min | 1,8 min | 0,78 min | 1,76 min | 0,55 min | **22,56 min** | **2,14 min** |
| IBD dataset | 1359 | 82,05 min | 23,52 min | 9,13 min | 5,66 min | 2,75 min | 356,48 min | **93,93 min** | **385,66 min** |

## 4 Discussion and conclusions

We describe DiTaxa, a method implementing a new paradigm for host disease status prediction and biomarker detection from 16S rRNA amplicon data. The main distinction of this approach from existing methods is substituting standard OTU-clustering [49] or sequence-level analysis [18] by segmenting 16S rRNA reads into the most frequent variable-length subsequences of a dataset. The proposed sequence segmentation, called Nucleotide-pair Encoding, is an unsupervised approach inspired by Byte-pair Encoding, a data compression algorithm that recently became popular in deep natural language processing. The identified subsequences represent commonly occurring sequence portions, which we found to be distinctive for taxa at varying evolutionary distances and highly informative for predicting host disease phenotypes. We compared the performance of DiTaxa to the state-of-the-art in disease phenotype prediction and biomarker detection, using human 16S datasets from metagenomic samples of periodontal, rheumatoid arthritis, and inflammatory bowel diseases, as well as a synthetic benchmark dataset. DiTaxa identified 17 of 29 taxa with confirmed links to periodontitis (recall= 0.59), while the OTU-based approach could only detect 3 of 29 organisms (recall= 0.10). In addition, we show that for the rheumatoid arthritis dataset, machine-learning classifiers trained to predict host disease phenotypes based on the NPE representation substantially outperformed OTU features (macro-F1 =0.76 compared to 0.65) and performed competitively for Crohn’s disease and synthetic datasets. Taxa predicted by DiTaxa for samples from patients with untreated rheumatoid arthritis (new onset RA) versus healthy individuals had *Prevotella copri* as the most significantly ranked, which was confirmed based on shotgun metagenomic analysis and in mouse experiments [71], while the standard workflow only predicted the genus of this taxon as relevant [71]. Due to the alignment- and reference free nature, DiTaxa can efficiently run on large datasets. The full analysis of a large 16S rRNA dataset of 1359 samples required ≈1.5 hours, where the standard pipeline took ≈6.5 hours with the same number of cores (20 cores). Although on smaller datasets the conventional workflow was faster than DiTaxa, the run-time difference of less than 30 minutes for those settings is worth the performance gain in phenotype prediction and biomarker detection. The applications of NPE representation are not limited to 16S rRNA data and it can be also applied to shotgun metagenomics or any other biological sequences to infer intrinsic features from data, instead of using parameter-dependent representations. Taken together, DiTaxa seems to provide a better solution for biomarker and phenotype detection than OTU-based methods. It thus could contribute to a better understanding of the microbial organisms associated with microbiome-related diseases and the development of personalized diagnostics and therapy procedures.

## Acknowledgements

Fruitful discussions with Curtis Huttenhower, Hinrich Schütze, Szymon Szafranski, and Benjamin Roth are gratefully acknowledged. P.C.M. received funding from German Research Foundation (315980449).

## Footnotes

1 Available at: https://www.ncbi.nlm.nih.gov/bioproject/PRJEB13679

2 Available at https://qiita.ucsd.edu/study/description/1939

3 Downloaded from http://datadryad.org/resource/doi:10.5061/dryad.d41v4

4 http://www.drive5.com/usearch/

## References

1. Costello, E. K. et al. Bacterial community variation in human body habitats across space and time. Sci. (New York, N.Y.) 326, 1694–7 (2009). URL http://www.sciencemag.org/content/326/5960/1694.short/ http://dx.doi.org/10.1126/science.1177486/ http://www.ncbi.nlm.nih.gov/pubmed/19892944/ http://www.pubmedcentral.nih.gov/articlerender.fcgi?artid=PMC3602444. DOI 10.1126/science.1177486.

2. Eck, A. et al. Robust microbiota-based diagnostics for inflammatory bowel disease. J. Clin. Microbiol. 55, 1720–1732 (2017). DOI 10.1128/JCM.00162-17.

3. Duvallet, C., Gibbons, S. M., Gurry, T., Irizarry, R. A. & Alm, E. J. Meta-analysis of gut microbiome studies identifies disease-specific and shared responses. Nat. Commun. 8, 1784 (2017). URL http://www.nature.com/articles/s41467-017-01973-8. DOI 10.1038/s41467-017-01973-8.

4. Pollock, J., Glendinning, L., Wisedchanwet, T. & Watson, M. The madness of microbiome: Attempting to find consensus “best practice” for 16S microbiome studies. Appl. Environ. Microbiol. AEM.02627–17 (2018). URL http://aem.asm.org/lookup/doi/10.1128/AEM.02627-17. DOI 10.1128/AEM.02627-17.

5. Cottier, F. et al. Advantages of meta-total rna sequencing (metrs) over shotgun metagenomics and amplicon-based sequencing in the profiling of complex microbial communities. npj Biofilms Microbiomes 4, 2 (2018).

6. Ward, D. M., Weller, R. & Bateson, M. M. 16S rRNA sequences reveal numerous uncultured microorganisms in a natural community. Nat. 345, 63–65 (1990). URL http://www.nature.com/doifinder/10.1038/345063a0. DOI 10.1038/345063a0.0801.3609.

7. Rideout, J. R. et al. Subsampled open-reference clustering creates consistent, comprehensive OTU definitions and scales to billions of sequences. PeerJ 2, e545 (2014). URL https://peerj.com/articles/545. DOI 10.7717/peerj.545.

8. Li, W. & Godzik, A. Cd-hit: A fast program for clustering and comparing large sets of protein or nucleotide sequences. Bioinforma. 22, 1658–1659 (2006). DOI 10.1093/bioinformatics/btl158.

9. DeSantis, T. Z. et al. Greengenes, a chimera-checked 16S rRNA gene database and workbench compatible with ARB. Appl. Environ. Microbiol. 72, 5069–5072 (2006). DOI 10.1128/AEM.03006-05.

10. Schloss, P. D. et al. Introducing mothur: Open-source, platform-independent, community-supported software for describing and comparing microbial communities. Appl. Environ. Microbiol. 75, 7537–7541 (2009). DOI 10.1128/AEM.01541-09.

11. Caporaso, J. G. et al. QIIME allows analysis of high-throughput community sequencing data (2010). DOI 10.1038/nmeth.f.303.NIHMS150003.

12. Lawley, B. & Tannock, G. W. Analysis of 16S rRNA Gene Amplicon Sequences Using the QIIME Software Package, vol. 1537 (Springer, 2017). URL http://www.ncbi.nlm.nih.gov/pubmed/27924593/ http://link.springer.com/10.1007/978-1-4939-6685-1{_}9.

13. Edgar, R. C., Haas, B. J., Clemente, J. C., Quince, C. & Knight, R. UCHIME improves sensitivity and speed of chimera detection. Bioinforma. 27, 2194–2200 (2011). DOI 10.1093/bioinformatics/btr381.

14. Hildebrand, F., Tadeo, R., Voigt, A. Y., Bork, P. & Raes, J. Lotus: an efficient and user-friendly otu processing pipeline. Microbiome 2, 30 (2014).

15. Kunin, V., Engelbrektson, A., Ochman, H. & Hugenholtz, P. Wrinkles in the rare biosphere: pyrosequencing errors can lead to artificial inflation of diversity estimates. Environ. microbiology 12, 118–123 (2010).

16. Schmidt, T. S. B., Matias Rodrigues, J. F. & von Mering, C. Ecological Consistency of SSU rRNA-Based Operational Taxonomic Units at a Global Scale. PLoS Comput. Biol. 10 (2014). DOI 10.1371/journal.pcbi.1003594.

17. He, Y. et al. Erratum to: Stability of operational taxonomic units: an important but neglected property for analyzing microbial diversity. Microbiome 3 (2015). URL http://www.microbiomejournal.com/content/3/1/34. DOI 10.1186/s40168-015-0098-1.

18. Callahan, B. J. et al. Dada2: high-resolution sample inference from illumina amplicon data. Nat. methods 13, 581 (2016).

19. Amir, A. et al. Deblur rapidly resolves single-nucleotide community sequence patterns. MSystems 2, e00191–16 (2017).

20. Nearing, J. T., Douglas, G. M., Comeau, A. M. & Langille, M. G. Denoising the denoisers: An independent evaluation of microbiome sequence error-correction methods. PeerJ PrePrints (2018).

21. Asgari, E., Garakani, K., McHardy, A. C. & Mofrad, M. R. Micropheno: Predicting environments and host phenotypes from 16s rrna gene sequencing using a k-mer based representation of shallow sub-samples. Bioinforma. J. (In press) bioRxiv–255018 (2018).

22. Statnikov, A. et al. A comprehensive evaluation of multicategory classification methods for microbiomic data. Microbiome 1, 11 (2013). URL http://microbiomejournal.biomedcentral.com/articles/10.1186/2049-2618-1-11. DOI 10.1186/2049-2618-1-11.

23. Carrieri, A. P. et al. Host Phenotype Prediction from Differentially Abundant Microbes Using RoDEO, 27–41 (Springer International Publishing, Cham, 2017). URL https://doi.org/10.1007/978-3-319-67834-4_3.

24. Kuczynski, J. et al. Direct sequencing of the human microbiome readily reveals community differences. Genome Biol. 11, 210 (2010). URL https://doi.org/10.1186/gb-2010-11-5-210. DOI 10.1186/gb-2010-11-5-210.

25. Parks, D. H. & Beiko, R. G. Measures of phylogenetic differentiation provide robust and complementary insights into microbial communities. The ISME J. 7, 173–183 (2012). URL https://doi.org/10.1038/ismej.2012.88. DOI 10.1038/ismej.2012.88.

26. Ley, R. E. et al. Obesity alters gut microbial ecology. Proc. Natl. Acad. Sci. 102, 11070–11075 (2005). URL http://www.pnas.org/cgi/doi/10.1073/pnas.0504978102. DOI 10.1073/pnas.0504978102.304.

27. Ley, R., Turnbaugh, P., Klein, S. & Gordon, J. Microbial ecology: human gut microbes associated with obesity. Nat. 444, 1022–3 (2006). URL http://europepmc.org/abstract/MED/17183309. DOI 10.1038/nature4441021a.NIHMS150003.

28. Turnbaugh, P. J. et al. A core gut microbiome in obese and lean twins. Nat. 457, 480–484 (2008). URL https://doi.org/10.1038/nature07540. DOI 10.1038/nature07540.

29. Mimee, M., Tucker, A. C., Voigt, C. A. & Lu, T. K. Programming a Human Commensal Bacterium, Bacteroides thetaiotaomicron, to Sense and Respond to Stimuli in the Murine Gut Microbiota. Cell Syst. 1, 62–71 (2015). DOI 10.1016/j.cels.2015.06.001.15334406.

30. Sheth, R. U., Cabral, V., Chen, S. P. & Wang, H. H. Manipulating Bacterial Communities by in situ Microbiome Engineering (2016). DOI 10.1016/j.tig.2016.01.005.15334406.

31. de la Fuente-Núñez, C. & Lu, T. K. CRISPR-Cas9 technology: applications in genome engineering, development of sequence-specific antimicrobials, and future prospects. Integr. Biol. 9, 109–122 (2017). URL http://xlink.rsc.org/?DOI=C6IB00140H. DOI 10.1039/C6IB00140H.

32. Dao, M. C. et al. Akkermansia muciniphila and improved metabolic health during a dietary intervention in obesity: Relationship with gut microbiome richness and ecology. Gut 65, 426–436 (2016). DOI 10.1136/gutjnl-2014-308778.

33. Paulson, J. N., Colin Stine, O., Bravo, H. C. & Pop, M. Differential abundance analysis for microbial marker-gene surveys. Nat. Methods 10, 1200–1202 (2013). DOI 10.1038/nmeth.2658.NIHMS150003.

34. Segata, N. et al. Metagenomic biomarker discovery and explanation. Genome Biol. 12 (2011). DOI 10.1186/gb-2011-12-6-r60. Segata, Nicola,2011, Metagenomic.

35. Kruskal, W. H. & Wallis, W. A. Use of Ranks in One-Criterion Variance Analysis. J. Am. Stat. Assoc. 47, 583–621 (1952). DOI 10.1080/01621459.1952.10483441.NIHMS150003.

36. Parks, D. H., Tyson, G. W., Hugenholtz, P. & Beiko, R. G. STAMP: Statistical analysis of taxonomic and functional profiles. Bioinforma. 30, 3123–3124 (2014). DOI 10.1093/bioinformatics/btu494.

37. Gevers, D. et al. The treatment-naive microbiome in new-onset Crohn’s disease. Cell Host Microbe 15, 382–392 (2014). DOI 10.1016/j.chom.2014.02.005.

38. Scher, J. U. et al. Expansion of intestinal Prevotella copri correlates with enhanced susceptibility to arthritis. eLife 2013 (2013). DOI 10.7554/eLife.01202.001.

39. Jorth, P. et al. Metatranscriptomics of the human oral microbiome during health and disease. MBio 5, e01012–14 (2014).

40. Angly, F. E., Willner, D., Rohwer, F., Hugenholtz, P. & Tyson, G. W. Grinder: a versatile amplicon and shotgun sequence simulator. Nucleic acids research 40, e94–e94 (2012).

41. Gage, P. A new algorithm for data compression. The C Users J. 12, 23–38 (1994).

42. Shibata, Y. et al. Byte pair encoding: a text compression scheme that accelerates pattern matching. Tech. Rep. DOI-TR-161, Dep. Informatics, (1999). URL https://pdfs.semanticscholar.org/1e94/41bbad598e181896349757b82af42b6a6902.pdf.

43. Chen, L., Lu, S. & Ram, J. Compressed pattern matching in dna sequences. In Computational Systems Bioinformatics Conference, 2004. CSB 2004. Proceedings. 2004 IEEE, 62–68 (IEEE, 2004).

44. Sennrich, R., Haddow, B. & Birch, A. Neural machine translation of rare words with subword units. In Proceedings of the 54th Annual Meeting of the Association for Computational Linguistics, 1715–1725 (Association for Computational Linguistics, 2016). URL http://www.aclweb.org/anthology/P16-1162. DOI 10.18653/v1/P16-1162.

45. Kudo, T. Subword regularization: Improving neural network translation models with multiple subword candidates. arXiv preprint arXiv:1804.10959 (2018).

46. Taft, D. H. et al. Intestinal microbiota of preterm infants differ over time and between hospitals. Microbiome 2, 36 (2014).

47. Wang, L., Li, P., Tang, Z., Yan, X. & Feng, B. Structural modulation of the gut microbiota and the relationship with body weight: compared evaluation of liraglutide and saxagliptin treatment. Sci. reports 6, 33251 (2016).

48. Bindels, L. B. et al. Synbiotic approach restores intestinal homeostasis and prolongs survival in leukaemic mice with cachexia. The ISME journal 10, 1456 (2016).

49. Edgar, R. C. Uparse: highly accurate otu sequences from microbial amplicon reads. Nat. methods 10, 996 (2013).

50. Yoon, S.-H. et al. Introducing ezbiocloud: a taxonomically united database of 16s rrna gene sequences and whole-genome assemblies. Int. journal systematic evolutionary microbiology 67, 1613–1617 (2017).

51. Wang, Q., Garrity, G. M., Tiedje, J. M. & Cole, J. R. Naive bayesian classifier for rapid assignment of rrna sequences into the new bacterial taxonomy. Appl. environmental microbiology 73, 5261–5267 (2007).

52. Breiman, L. Random Forests. Mach. Learn. 45, 5–32 (2001).

53. Camacho, C. et al. Blast+: architecture and applications. BMC bioinformatics 10, 421 (2009).

54. Kullback, S. & Leibler, R. A. On information and sufficiency. The annals mathematical statistics 22, 79–86 (1951).

55. Segata, N., Börnigen, D., Morgan, X. C. & Huttenhower, C. Phylophlan is a new method for improved phylogenetic and taxonomic placement of microbes. Nat. communications 4, 2304 (2013).

56. Polak, D. et al. Mouse model of experimental periodontitis induced by Porphyromonas gingivalis Fusobacterium nucleatum infection: Bone loss and host response. J. Clin. Periodontol. 36, 406–410 (2009). DOI 10.1111/j.1600-051X.2009.01393.x.

57. Kimura, S. et al. Induction of experimental periodontitis in mice with Porphyromonas gingivalis-adhered ligatures. J. Of Periodontol. 71, 1167–1173 (2000). URL http://eutils.ncbi.nlm.nih.gov/entrez/eutils/elink.fcgi?dbfrom=pubmed{&}id=10960025{&}retmode=ref{&}cmd=prlinks{%}5Cnpapers2://publication/doi/10.1902/jop.2000.71.7.1167. DOI 10.1902/jop.2000.71.7.1167.

58. Hajishengallis, G. et al. Low-abundance biofilm species orchestrates inflammatory periodontal disease through the commensal microbiota and complement. Cell host & microbe 10, 497–506 (2011).

59. Perez-Chaparro, P. et al. Newly identified pathogens associated with periodontitis: a systematic review. J. dental research 93, 846–858 (2014).

60. Liu, B. et al. Deep sequencing of the oral microbiome reveals signatures of periodontal disease. PLoS ONE 7 (2012). DOI 10.1371/journal.pone.0037919.

61. Scher, J. U. et al. Periodontal disease and the oral microbiota in new-onset rheumatoid arthritis. Arthritis Rheum. 64, 3083–3094 (2012). DOI 10.1002/art.34539.NIHMS150003.

62. Oz, H. S. & Puleo, D. a. Animal models for periodontal disease. J. biomedicine & biotechnology 2011, 754857 (2011). URL http://www.pubmedcentral.nih.gov/articlerender.fcgi?artid=3038839{&}tool=pmcentrez{&}rendertype=abstract. DOI 10.1155/2011/754857.

63. Teles, R., Teles, F., Frias-Lopez, J., Paster, B. & Haffajee, A. Lessons learned and unlearned in periodontal microbiology. Periodontol. 2000 62, 95–162 (2013).

64. Socransky, S., Haffajee, A., Cugini, M., Smith, C. & Kent, R. Microbial complexes in subgingival plaque. J. clinical periodontology 25, 134–144 (1998).

65. Schlafer, S. et al. Filifactor alocis-involvement in periodontal biofilms. BMC microbiology 10, 66 (2010).

66. Schillinger, C. et al. Co-localized or randomly distributed? pair cross correlation of in vivo grown subgingival biofilm bacteria quantified by digital image analysis. PLoS One 7, e37583 (2012).

67. Aas, J. A., Paster, B. J., Stokes, L. N., Olsen, I. & Dewhirst, F. E. Defining the normal bacterial flora of the oral cavity. J. clinical microbiology 43, 5721–5732 (2005).

68. Ximénez-Fyvie, L. A., Haffajee, A. D. & Socransky, S. S. Microbial composition of supra-and subgingival plaque in subjects with adult periodontitis. J. clinical periodontology 27, 722–732 (2000).

69. Abusleme, L. et al. The subgingival microbiome in health and periodontitis and its relationship with community biomass and inflammation. The ISME journal 7, 1016 (2013).

70. Ledder, R. G. et al. Molecular analysis of the subgingival microbiota in health and disease. Appl. environmental microbiology 73, 516–523 (2007).

71. Scher, J. U. et al. Expansion of intestinal prevotella copri correlates with enhanced susceptibility to arthritis. elife 2 (2013).

